# Differential proteomic analysis by SWATH-MS unravels the most dominant mechanisms underlying yeast adaptation to non-optimal temperatures under anaerobic conditions

**DOI:** 10.1101/2020.01.06.895581

**Authors:** Tânia Pinheiro, Ka Ying Florence Lip, Estéfani García-Ríos, Amparo Querol, José Teixeira, Walter van Gulik, José Manuel Guillamón, Lucília Domingues

## Abstract

Elucidation of temperature tolerance mechanisms in yeast is essential for enhancing cellular robustness of strains, providing more economically and sustainable processes. We investigated the differential responses of three distinct *Saccharomyces cerevisiae* strains, an industrial wine strain, ADY5, a laboratory strain, CEN.PK113-7D and an industrial bioethanol strain, Ethanol Red, grown at sub- and supra-optimal temperatures under chemostat conditions. We employed anaerobic conditions, mimicking the industrial processes. The proteomic profile of these strains was performed by SWATH-MS, allowing the quantification of 997 proteins, data available via ProteomeXchange (PXD016567). Our analysis demonstrated that temperature responses differ between the strains; however, we also found some common responsive proteins, revealing that the response to temperature involves general stress and specific mechanisms. Overall, sub-optimal temperature conditions involved a higher remodeling of the proteome. The proteomic data evidenced that the cold response involves strong repression of translation-related proteins as well as induction of amino acid metabolism, together with components related to protein folding and degradation while, the high temperature response mainly recruits amino acid metabolism. Our study provides a global and thorough insight into how growth temperature affects the yeast proteome, which can be a step forward in the comprehension and improvement of yeast thermotolerance.

## Introduction

The yeast *Saccharomyces cerevisiae* is used as a microbial cell factory in a wide range of industrial applications, from the production of beer, wine, cider and bread to biofuels and pharmaceuticals^1^. However, these industrial processes impose severe stresses that impact the efficiency of this cell factory. Among these factors, temperature is one of the most relevant variables, with a direct influence on yeast growth and fermentation performance, representing a major economic concern in the biotechnology industry. In fact, the industry spends large amounts of energy in heating or cooling and, in many cases, the optimum growth temperature is sacrificed^2^. Hence, a significant cost reduction and productivity increase would be achieved in fermentations with better-adapted yeast to ferment at non-optimal temperatures. For these reasons, there is a clear interest in identifying the mechanisms that underlie yeast adaptation at high and low temperatures, as this would make it possible to improve or engineer yeast strains with enhanced robustness.

In past years, some attempts have been made to elucidate the thermal response of *S. cerevisiae*^3–11^. Much of the current knowledge has been derived from studies on the effects of abrupt temperature shocks rather than prolonged thermal stress. However, the magnitude of the stress response is highly influenced by the rate of change of environmental conditions, which is why it is crucial to distinguish between transient stress responses and stress adaptation^12^. Studies on transient stress responses are typically conducted in batch cultures, whereby stress in the form of a shock is applied, so that cells are more prone to triggering fast and highly dynamic stress response phenomena^13^. On the other hand, prolonged exposure to nonlethal stimuli leads to acclimation; i.e., establishment of a physiological state in which regulatory mechanisms, like gene expression, fully adapted to suboptimal environmental conditions^13^. This type of studies is carried out in continuous culture experiments, such as chemostats, and has the advantage of discriminating the effects of the growth rate and the applied stressor^14^. Moreover, most of these approaches have used genome-wide transcriptomic data^3,7,9,15^. An inspection of the transcriptome provides accurate identification of the genes which are active in cells. Nevertheless, strong gene expression and high mRNA levels, does not necessarily mean that the corresponding protein is also abundant or indeed active in the cell. Although proteomic analysis techniques are not as straightforward as those used in transcriptomics, they offer the advantage of studying proteins, which represent the actual functional molecules in the cell^16^. To our knowledge, there are only two proteomic studies that examine the tolerance mechanisms of yeast grown at high temperature^4,5^, and one at low temperature^17^. However, only the study performed by García-Ríos et al. (2016) was conducted under adapted stress conditions such as chemostats^17^.

Yeast thermotolerance is recognized to be a complicated quantitative trait. In addition, the heterogeneity among *S. cerevisiae* strains is significant, with laboratory strains presenting lower thermotolerance than robust industrial and natural yeast strains^18,19^. Still, studies are generally focused on only one background strain, which is not enough to elucidate the intricate mechanism of thermotolerance.

The goal of the present study is to make significant advances in our understanding of yeast response to sub- and supra-optimal temperatures. To this end, we used SWATH-MS (sequential window acquisition of all theoretical spectra-mass spectrometry), a highly accurate quantification technique, to characterize the proteome remodeling of three phenotypically distinct *S. cerevisiae* strains Ethanol Red (bioethanol strain), – a previously selected high temperature tolerant strain^20^, ADY5 (wine strain), a previously selected low temperature tolerant strain^20^, and CEN.PK113-7D (laboratory reference strain). To eliminate interference by specific growth rate, the strains were grown at 12 °C, 30 °C and 39 °C, in anaerobic chemostat cultures, at a fixed specific growth rate of 0.03 h^-1^, enabling an accurate investigation of the impact of temperature. Anaerobic conditions were chosen to mimic the process of industrial fermentation of alcoholic beverages and bioethanol production. Furthermore, it also prevents the possible effect of temperature-dependent oxygen solubility and of temperature dependent distribution of sugar metabolism over respiration and alcoholic fermentation^13^. Using independent culture replicates and stringent statistical filtering, we defined a robust dataset of 259 differential quantified proteins. Beyond uncovering the most dominant mechanisms underlying temperature adaptation, the use of three strains with different degrees of thermotolerance represents a comprehensive strategy that will allow the identification of key strain-dependent and temperature-dependent physiological differences, providing the necessary knowledge for the further production of tailored thermotolerant yeasts.

## Materials and Methods

### Yeast strains, cultivation media and inoculum preparation

The yeast strains used in this work were *S. cerevisiae* ADY5 (Lallemand Inc., Canada), a commercial wine strain, *S. cerevisiae* Ethanol Red^®^ (Fermentis, S.I. Lesaffre, France), a commercial bioethanol strain, and the haploid laboratory strain *S. cerevisiae* CEN.PK113-7D (Fungal Biodiversity Centre, Utrecht, The Netherlands). The ADY5 and Ethanol Red were selected from a screening involving 12 industrial strains, as the most tolerant to 12 °C and 39 °C^20^, respectively.

Working stocks were prepared by cultivation in YPD medium, containing per L: 10 g Bacto yeast extract, 20 g Bacto peptone and 20 g D-glucose. After addition of 30 % (v/v) glycerol, culture aliquots were stored in sterilized Eppendorf tubes at −80°C.

Inocula for the chemostat cultivations were grown aerobically at 220 *rpm* at 30 °C in 2 L Erlenmeyer flasks containing 400 mL of filter sterilized medium containing per L: 5 g (NH_4_)_2_SO_4_, 3 g KH_2_PO_4_, 0.5 g MgSO_4_·_7_H_2_O, 15 g glucose. H_2_O, 1.0 mL of trace element solution, and 1.0 mL vitamin solution. Trace element and vitamin solutions were prepared as described by Verduyn et al. (1992)^21^. The preculture medium was filter sterilized using a 0.2 μm membrane filter (Supor AcroPak 20, Pall, Port Washington, USA).

The medium for anaerobic chemostat cultivation as well as the preceding batch cultivations contained per L: 5.0 g (NH_4_)_2_SO_4_, 3.0 g KH_2_PO_4_, 0.5 g·MgSO_4_·7H_2_O, 22.0 g D-glucose ·H_2_O, 0.4 g Tween80, 10 mg ergosterol, 0.26 g antifoam C (Sigma-Aldrich, Missouri, USA), 1.0 mL trace element solution, and 1.0 mL vitamin solution. The medium was filter sterilized using a 0.2 μm Sartopore 2 filter unit (Sartorius Stedim, Goettingen, Germany).

### Chemostat cultivations

All chemostat cultivations were carried out at a dilution rate of 0.030 ± 0.002 h^-1^ in 7 L bioreactors (Applikon, Delft, The Netherlands) equipped with a DCU3 control system and MFCS data acquisition and control software (Sartorius Stedim Biotech, Goettingen, Germany). The reactor vessels were equipped with norprene tubing, to minimize the diffusion of oxygen into the vessels, and were sterilized by autoclaving at 121 °C.

During chemostat operation, the sterile feed medium was pumped into the reactor vessel at a constant flowrate using a peristaltic pump (Masterflex, Barrington, USA), such that the outflow rate of the culture broth was 120 ± 1 g·h^-1^. The broth mass in the reactor was maintained at 4.00 ± 0.05 kg, by discontinuous removal of culture into a sterile effluent vessel, via a pneumatically operated valve in the bottom of the reactor and a peristaltic pump, which were operated by weight control. Therefore, the complete reactor was placed on a load cell (Mettler Toledo, Tiel, The Netherlands). Also, the effluent vessel was placed on a load cell of which the signal was continuously logged for accurate determination of the dilution rate of the chemostat and manual adjustment of the medium feed rate if needed.

The cultivations were carried out at temperatures of either 12.0 ± 0.1 °C, 30.0 ± 0.1°C or 39.0 ± 0.1 °C, by pumping cooled or heated water through the stainless-steel jacket surrounding the bottom part of the reactor vessel, using a cryothermostat (Lauda RE630, Lauda-Königshofen, Germany). The water temperature of the cryothermostat was controlled by using the signal of a Pt 100 temperature sensor inside the reactor, for accurate measurement and control of the cultivation temperature. Anaerobic conditions were maintained by continuously gassing of the reactor with nitrogen gas at a flowrate of 1 SLM (standard liter per minute) using a mass flow controller (Brooks, Hatfield, USA). Also, the feed medium was kept anaerobic by sparging with nitrogen gas. The nitrogen gas was sterilized by passing through sterile hydrophobic plate filters with a pore size of 0.2 μm (Millex, Millipore, Billerica, USA). The culture broth in the reactor was mixed using one 6-bladed Rushton turbine (diameter 80 mm) operated at a rotation speed of 450 rpm. The pH was controlled at 5.00 ± 0.05 by automatic titration with 4M KOH. The bioreactor was inoculated with 400 mL of pre-culture and subsequently operated in batch-mode, allowing the cells to grow at the same temperature as the chemostat culture and to achieve enough biomass at the start of the chemostat phase. The exhaust gas from the chemostat was passed through a condenser kept at 4 °C and then through a Perma Pure Dryer (Inacom Instruments, Overberg, The Netherlands) to remove all water vapor and subsequently entered a Rosemount NGA 2000 gas analyzer (Minnesota, USA) for measurement of the CO_2_ concentration. When the CO_2_ level of the exhaust gas during the batch cultivation dropped significantly, close to the level after the preculture inoculation, this indicated the end of the batch phase. Thereafter the culture was switched to chemostat mode. Sampling was carried out during steady state conditions, after stable values of the CO_2_ level in the exhaust gas and the biomass dry weight concentration were obtained. Triplicate samples were taken from each chemostat cultivation approximately every 48 h during the steady state for measurement of cell dry weight and proteomics analysis. The steady-state of the chemostat cultures was confirmed by the stable CO_2_ off gas profile and steady dry-weight measurements^20^.

Biomass dry weight was determined by filtration of 5 g of chemostat broth over pre-dried nitrocellulose filters (0.45 μm pore size, Gelman laboratory, Ann Arbor, USA), which were wetted with 1 mL demineralized water before filtration. After filtration of the broth, 10 mL of demineralized water was used to wash the cells on the filters, which were subsequently dried in an oven at 70°C for two days. Before weighing, the filters were allowed to cool down in a desiccator for two hours.

For proteomics analysis 3 x10 g of chemostat broth was immediately cooled down to around 4°C after sampling by pouring into a tube containing cold steel beads. After cold centrifugation the cell pellets were snap frozen in liquid nitrogen and stored at −80°C until analysis.

### Protein extraction and digestion

Cell pellets were then resuspended in 500 μL of UTC lysis buffer, containing 8 M urea, 2 M thiourea and 4% 3-[(3-cholamidopropyl)dimethylammonio]-1-propanesulfonate, (CHAPS), followed by vigorous stirring at 5 °C during 1 h. After, the samples were treated with 10 % (final concentration) of trichloroacetic acid (TCA) and incubated at 4 °C overnight. Protein extracts from total cell were clarified by centrifugation at 15000*g* during 10 min and precipitated according to the TCA/Acetone protocol. Briefly, the treated samples were diluted (equal volume of the initial sample) in cold acetone solution, stirred and stored at 4 °C during 10 min. Then, the samples were subjected to centrifugation 15000*g* for 15min and the supernatants were discarded. The total of the precipitated proteins was taken for one dimensional sodium dodecyl sulfatepolyacrylamide gel electrophoresis (1D SDS-PAGE). The samples were loaded but not resolved.

The protein samples were digested with 500 ng of sequencing grade trypsin (V511, Promega Co., Madison, WI, USA) and incubated at 37 °C following the protocol described by Shevchenko et al.^22^. The trypsin digestion was stopped by the addition of 10 % trifuoroacetic (TFA) and the supernatant, containing the non-extracted digests, was removed, leaving behind the sliced gels in the Eppendorf tube, which were dehydrate with pure acetonitrile (ACN). The new peptide solutions were carefully combined with the corresponding supernatant and dried in a speed vacuum. Next, they were re-suspended in 2 % ACN and 0.1 % TFA prior to liquid chromatography and tandem mass spectrometry (LC–MS/MS) analyses. The volume was adjusted according to the intensity of the staining.

### LC-MS/MS analyses

#### Spectral library building

Liquid chromatography and tandem mass spectrometry (LC-MS/MS): 5 μL of a pool of all the digested samples were loaded into a trap column (Nano LC Column, 3μ C18 CL, 75 μm x 15 cm; Eksigent Technologies, Dublin, CA, USA) and desalted with 0.1 % TFA at 3μL/min during 5 min. The peptides were loaded into an analytical column (LC Column, 3 μm particles size C18-CL, 75 μm ×12cm, Nikkyo Technos Co^®^, Tokyo, Japan), equilibrated in 5 % ACN and 0.1 % formic acid (FA). The peptide elution was carried out with a linear gradient of 5 % to 35 % buffer B (B: ACN, 0.1 % FA) in A (A: 0.1 % FA) at a constant flow rate of 300 nL/min over 240 min. The eluted peptides were analyzed in a mass spectrometer nanoESI-qQTOF (5600 TripleTOF, AB SCIEX). The TripleTOF was operated in information-dependent acquisition mode, in which a 250-ms TOF MS scan from 350–1250 m/z, was performed, followed by 150-ms product ion scans from 350–1500 m/z on the 25 most intense 2–5 charged ions. The rolling collision energies equations were set for all ions as for 2+ ions according to the equation |CE|=(slope) × (m/z) + (intercept), with Slope = 0.0575 and Intercept = 9.

#### SWATH acquisition

The digested peptides recovered were examined by LC using a NanoLC Ultra 1-D plus Eksigent (Eksigent Technologies, Dublin, CA, USA) connected to a TripleTOF 5600 mass spectrometer (AB SCIEX, Framingham, MA, USA). Briefly, 5 μL of each digested sample were loaded onto a trap Nano LC pre-column (3 μm particles size C18-CL, 0.5 mm × 300 μm, Eksigent Technologies) and desalted with 0.1 % TFA at 3μL/min during 5 min. Then, the digested peptides present in the samples were separated using an analytical column (LC Column, 3 μm particles size C18-CL, 75 μm ×12cm, Nikkyo Technos Co, Tokyo, Japan), equilibrated in 5 % ACN and 0.1 % FA. The peptide elution was carried out with a linear gradient of 5% to 35% of ACN containing 0.1% FA at a constant flow rate of 300 nL/min over 90 min. The eluted peptides were thereafter analyzed with a spectrometer nanoESI-qQTOF (5600 TripleTOF, AB SCIEX) coupled to the NanoLC sysntem. The TripleTOF was operated in SWATH mode, in which a 0.050s TOF MS scan from 350 to 1250m/z was performed, followed by 0.080s product ion scans from 350 to 1250 m/z in the 32 defined windows (3.05 sec/cycle). The rolling collision energies equations were set for all ions as for 2+ ions according to the equation |CE|=(slope) × (m/z) + (intercept), with Charge = 2; Slope = 0.0575 and Intercept = 9.

### Protein identification and quantification

For spectral library building after LC-MS/MS analysis, the generated 5600 TripleTof.wiff data-files were processed using ProteinPilot search engine (version 5.0, AB SCIEX). The Paragon algorithm of ProteinPilot^23^ was used to search against the Uniprot database (SwissProt-032018.fasta, 556819 proteins in database) with the following parameters: trypsin specificity, cys-alkylation (IAM), no taxonomy restricted, and the search effort set to through and FDR correction for proteins. The proteins identified were grouped through the Protein-Pilot Pro ™ group algorithm exclusively from observed peptides and based on MS / MS spectra. Then, unobserved regions of protein sequence play no role for explaining the data.

The resulting Protein-Pilot group file was used by PeakView® (version 2.1, AB SCIEX) to quantify proteins, by chromatographic area, in SWATH runs. For every protein, in the spectral library a maximum of 50 peptides were quantified among those with a confidence threshold of 95 % and a false discovery rate (FDR) lower than 1 %. Shared peptides were also excluded. For every peptide a maximum of 6 transitions (fragment ions) were quantified. The peptide retention times were normalized among samples using high confident peptides of main proteins.

### Statistical analysis

The identified and quantified peak intensities of all the conditions (temperature and strains) were first transformed by the logarithm with base 2. Further, in order to filter the most differentiable proteins, an elastic-net penalized logistic regression (ENLR) model was performed to obtain the estimation of the coefficients. We use the glmnet package of R software to adjust the regression model^24^ and with the train function of the caret package to obtain a possible value for α and λ. The train function of the caret package was used to obtain the values for parameters needed for the regression model with Elastic net penalty. Train function sets up a grid of adjustment parameters for a number of classification and regression routines, adjusts to each model and calculates an efficiency measurement based on resampling. Within this function, tuneGrid parameter was used with an array of data with possible adjustment values. This matrix has a range of values that goes from the parameters that the glmnet function takes when alpha = 0 to alpha = 1. In this way the train function gives us a possible value for alpha and another for lambda. For each of the comparisons, we have applied a regression model with Elastic Net penalty, so we have had different parameters for each case.

The explanatory capacity of resulting selected proteins was shown using heatmaps after z-score normalization. To validate this methodology, multiple comparisons were performed by applying a False Discovery Rate (FDR). We also performed a PLS-DA classification of the proteins by using the mixOmics packet of R software. VIP function shows the importance of each of the explanatory variables in the projection which allow us to determine which of the variables is more important to predict the response variable. When vip value is greater than 1.5, the influence of the explanatory variable on the response variable is very high. We have checked that the proteins obtained with the Elastic Net regression model presented vip greater than 1.

Venn diagrams of the proteins with different concentrations at low temperature and high temperature were created using Venny online software (Venny, http://bioinfogp.cnb.csic.es/tools/venny/index.html), to select the common proteins from among the analyses. Principal component analysis (PCA) was performed using StatSoft, Inc. (2004) STATISTICA, version 7.0 (www.statsoft.com).

### Functional annotation and enrichment analysis

To explore whether certain biological processes are enriched among the proteins that were found significantly changed with the temperature, a gene enrichment analysis was manually performed using the comprehensive bioinformatics tool for functional annotation Swiss-Prot/TrEMBL (http://www.uniprot.org/uniprot) and evolutionary genealogy of genes: NOGs (http://eggnog.embl.de) databases.

## Results and discussion

### Overview of quantitative proteomic analysis

Our study presents the first proteomic characterization of three *Saccharomyces* strains grown at optimal temperature (i.e., 30 °C) versus low (i.e., 12 °C) and high temperature (i.e., 39 °C). This strategy allowed us to assess both the extent of the changes in protein expression levels in response to temperature and the degree of variance in the proteome expression profiles in yeasts from different ecological niches.

Quantitative proteomic data were obtained using the SWATH mass spectrometry (SWATH-MS) technology and resulted in the quantification of 997 unique proteins for all the samples, that is, the three strains cultivated at three temperatures with three biological replicates for each condition (27 samples in total). The full set of protein quantification data is provided in supplemental Table S1. To uncover the proteins significantly associated with temperature response, we employed elastic net regression analysis^25^, which is a method that allows the selection of highly correlated groups of variables. From the 997 proteins detected, the elastic net was able to significantly reduce the dataset in all the conditions, as can be seen in Table 1. Only these 259 selected proteins were considered in the following analysis.

To obtain an overview of the proteomic variability between the three strains, a Principal Component Analysis (PCA) was carried out (Figure 1).

**Figure 1 –.**
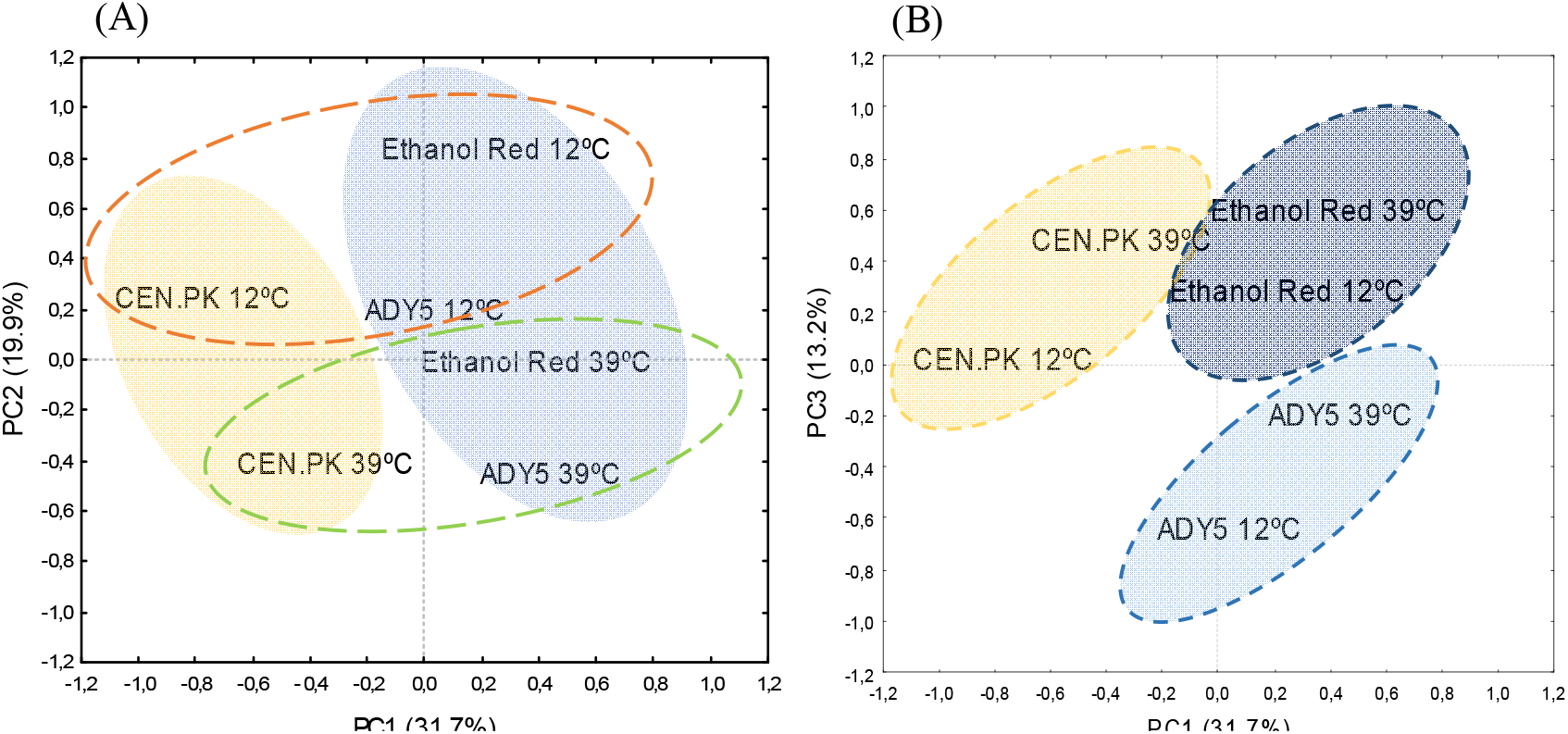
Principal component analysis (PCA) score plot from fold-change values of the differential expressed proteins in Ethanol Red, ADY5 and CEN.PK at 12 °C and 39 °C. (A) PCA Loadings of the second and first principal components (PC1 *vs* PC2); (B) PCA Loadings of the third and first principal components (PC1 *vs* PC3). The explained variances are shown in brackets. To make clear the hidden patterns in the data set, the different groups discriminated by each of the components (PC1, PC2 and PC3) were highlighted using colored ellipses and dashed lines.

The PCA score plot of the first three principal components accounted for 64.8 % of the total variance. The use of these components in a 2D representation mainly allowed the effective separation of the samples based on the strain and growth temperature. The first principal component (PC1) captured the largest variance of the data (31.7 %) and evidenced inter-strain differences, with a clear differentiation of the laboratory strain CEN.PK from the two industrial strains under study. The second PCA axis (PC2) accounted for 19.9 % of the total variance and clearly separated the strains according to the growth temperature. Thus, our preliminary proteome comparisons reveal that there are substantial differences at the molecular level in how the three strains respond to temperature changes and that the response to high and low temperature is quite distinct. Still, the third component (PC3) suggest that there are some general mechanisms of temperature response in each yeast strain, which probably corresponds to general stress response proteins.

### Biological mechanisms that govern cellular response to sub- and supra-physiological temperatures

The ability to adapt to non-optimal temperatures is potentially accompanied by changes in protein expression. To examine these changes, we evaluated the differentially expressed proteins of CEN.PK, Ethanol Red and ADY5 at 12 °C and 39 °C, compared to the respective control temperature (30 °C). Gene ontology analysis was performed to determine the main functional categories arising from the up- and down- regulated proteins. A total of 19 enriched GO terms (Table 1) were identified revealing the complexity of the cell’s response to temperature changes. A significant amount of proteins with unknown function were also identified.

The distribution of the differentially regulated proteins in functional categories differed considerably between the two temperature conditions. In fact, in all strains the number of differentially expressed proteins during cultivation at low temperature was much higher than at high temperature, indicating greater remodeling of the proteome. More specifically, in the ADY5 strain, the number of differentially regulated proteins at 12 °C was 85, whereas at 39 °C was 54; in the case of Ethanol Red, we identified 54 proteins at low temperature and 37 at high temperature; in CEN.PK, the discrepancy was even higher, with 104 differentially regulated proteins at low temperature, and only 24 at high temperature.

**Table 1 –.** Functional classification of differentially regulated protein expressed in ADY5, CEN.PK and Ethanol Red at 12 °C and 39 °C.

| Functional category | 12°C |  |  |  |  |  | 39°C |  |  |  |  |  |
| --- | --- | --- | --- | --- | --- | --- | --- | --- | --- | --- | --- | --- |
|  | A-D<sup>a</sup> | A-U<sup>a</sup> | C-D<sup>a</sup> | C-U<sup>a</sup> | E-D<sup>a</sup> | E-U<sup>a</sup> | A-D<sup>a</sup> | A-U<sup>a</sup> | C-D<sup>a</sup> | C-U<sup>a</sup> | E-D<sup>a</sup> | E-U<sup>a</sup> |
| Protein folding and degradation | 6 | 10 | 4 | 8 | 4 | 2 | 2 | 14 | 2 | 4 | 1 | 5 |
| Amino acid metabolism | 2 | 3 | 1 | 8 | 2 | 9 | 5 | 2 | 4 | 1 | 4 | 2 |
| Coenzyme metabolism | 2 | 0 | 1 | 2 | 1 | 0 | 0 | 0 | 1 | 0 | 0 | 0 |
| Inorganic ion metabolism | 2 | 0 | 0 | 2 | 0 | 0 | 0 | 0 | 1 | 0 | 0 | 0 |
| Signal transduction | 1 | 0 | 1 | 0 | 1 | 0 | 0 | 0 | 1 | 0 | 0 | 1 |
| Energy metabolism | 2 | 5 | 1 | 7 | 0 | 5 | 2 | 0 | 1 | 0 | 0 | 1 |
| Translation | 12 | 1 | 17 | 2 | 5 | 0 | 0 | 3 | 0 | 4 | 0 | 4 |
| Intracellular trafficking, secretion, and vesicular transport | 2 | 1 | 0 | 2 | 0 | 0 | 0 | 2 | 0 | 1 | 0 | 0 |
| Nucleotide metabolism | 2 | 1 | 2 | 4 | 0 | 1 | 2 | 3 | 0 | 1 | 1 | 4 |
| Carbohydrate metabolism | 0 | 6 | 2 | 8 | 3 | 3 | 1 | 2 | 0 | 0 | 2 | 1 |
| Cell cycle control, cell division, chromosome partitioning | 0 | 1 | 0 | 1 | 0 | 0 | 0 | 0 | 0 | 0 | 0 | 0 |
| Transcription | 3 | 0 | 1 | 2 | 1 | 1 | 0 | 2 | 0 | 0 | 0 | 2 |
| Replication, recombination and repair | 2 | 0 | 1 | 1 | 1 | 0 | 0 | 0 | 0 | 0 | 0 | 0 |
| Lipid metabolism | 0 | 0 | 2 | 1 | 5 | 1 | 0 | 0 | 0 | 0 | 0 | 1 |
| Secondary metabolites metabolism | 0 | 0 | 0 | 1 | 0 | 0 | 0 | 0 | 0 | 0 | 0 | 0 |
| Cytoskeleton | 1 | 0 | 0 | 0 | 0 | 0 | 0 | 1 | 0 | 0 | 0 | 0 |
| Chromatin structure and dynamics | 1 | 0 | 0 | 0 | 0 | 0 | 0 | 0 | 0 | 0 | 0 | 0 |
| RNA processing and modification | 3 | 0 | 0 | 0 | 0 | 0 | 0 | 1 | 0 | 0 | 0 | 1 |
| Cell wall biogenesis | 0 | 0 | 0 | 0 | 2 | 0 | 0 | 0 | 0 | 0 | 0 | 1 |
| Unknown Function | 8 | 8 | 2 | 20 | 4 | 3 | 1 | 11 | 1 | 2 | 2 | 3 |
| Total | 49 | 36 | 35 | 69 | 29 | 25 | 13 | 41 | 11 | 13 | 10 | 27 |

<sup>a</sup> The number of up (U) – or down (D)- regulated proteins in ADY5 (A), CEN.PK (C) and Ethanol Red (E) strains.
Scale bar: 1-5 6-10 11-15 16-20

Regarding the overall response to high and low temperatures, we found some resemblances because similar functions are induced or repressed under both experimental settings. The most remarkable groups of regulated proteins fell into the translation category, along with protein folding and degradation, and amino acid metabolism. In addition, at 12 °C, the transport and metabolism of carbohydrates as well as the production and conversion of energy also seem to be important categories. It is worth mentioning that at 12 °C there was a strong downregulation of proteins related to protein synthesis (translation process), particularly in the ADY5 and CEN.PK strains. Furthermore, the low temperature response of these two strains was also characterized by an increase in the levels of several proteins related to folding and degradation processes. The specific role of these functions and respective proteins will be explored in the next sections.

### Strain-specific responses to sub- and supra-optimal temperature

We further compared the proteome profile of the same strain but at different temperatures (12/30 °C versus 39/30 °C) to evaluate a possible stress-specific response. Regarding CEN.PK the comparison between the two temperature conditions showed that 92 and 12 proteins were uniquely differentially expressed at low and high temperature, respectively, whereby 12 proteins were differentially expressed at both conditions (Figure 2A). The shared proteins are mostly associated with translation, amino acid transport and protein folding and degradation.

**Figure 2 –.**
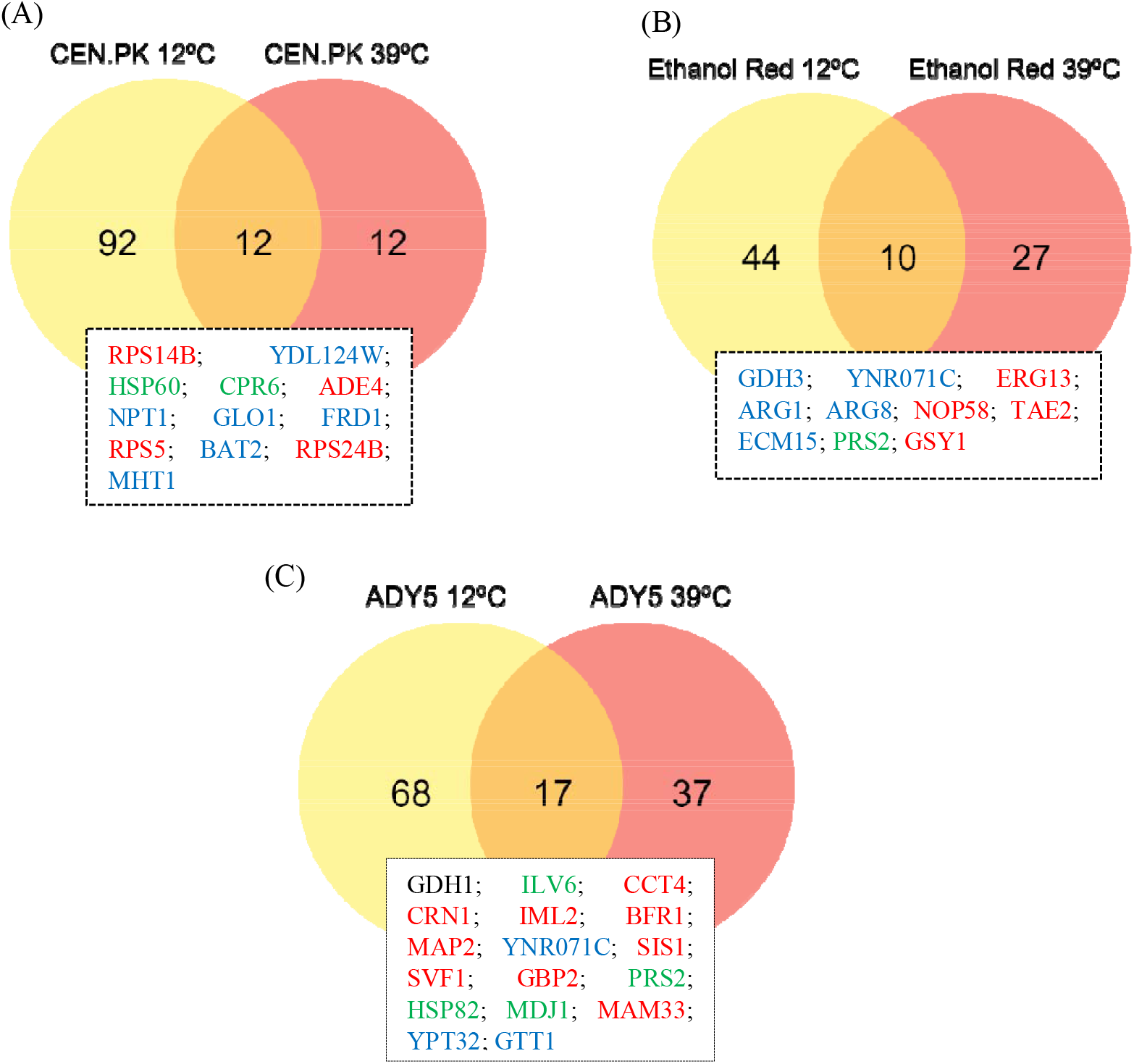
Venn diagram indicating the overlap of differentially expressed proteins between 12 °C and 39 °C conditions, compared to the respective control temperature (30 °C), in CEN.PK (A), Ethanol Red (B) and ADY5 (C). The standard name of the proteins common to both temperature settings are provided in the text boxes. Proteins are colored according to significant change as follows: red (increased at 39 °C and decreased at 12 °C), blue (decreased at 39 °C and increased at 12 °C), green (increased at both temperatures), or black (decreased at both temperatures).

In the case of Ethanol Red, it was observed that of the 81 identified proteins, 44 were unique to the cold response, 27 were unique to the heat response, and 10 were common to both (Figure 2B). Among the group of common proteins, we found proteins that have widespread functions in amino acid metabolism, carbohydrate metabolism, lipid metabolism, transcription, nucleotide metabolism and cell wall biogenesis.

In the ADY5 experiments we detected a similar behavior, with 17 proteins being involved in both high and low temperature response (Figure 2C). This group comprises proteins involved in several cellular functions such as amino acid metabolism (Gdh1, Ilv6), protein folding and degradation (Cct4, Sis1, Hsp82, Mdj1, Gtt1), RNA processing and modification (Svf1), translation (Map2), carbohydrate metabolism (Ynr071c), nucleotide metabolism (Prs2) and proteins with unknown functions (Crn1, Iml2, Bfr1, Gbp2, Mam33, Ypt32).

It should be noted that the proteins identified as common between low and high temperature response in most cases displayed opposite trends, which means that the proteins induced at 12 °C were repressed at 39 °C and vice versa. In a previous study on the molecular response to temperature, Strassburg et al. (2010) noticed that high and low temperatures cause distinct changes at both the metabolic and the transcript level^3^. They proposed that the yeast’s response to temperature involves two phases. First, there is the general stress response, which, through the perception of a stress-induced signal, allows the system to cope with cell damage and death; and second, there are specific adaptive responses to prolonged high and low temperature stress. Comparing the profile of some genes obtained at heat and cold conditions, they observed an opposite trend of regulation, in agreement with our results. Notwithstanding, a minor group of proteins was found to be induced or repressed in both temperature regimes. Specifically, for CEN.PK, Hsp60 and Cpr6 are upregulated at 12 °C and 39 °C, compare to the control temperature. These results evidenced that, in this strain, non-optimal temperature conditions impair protein folding processes.

In relation to Ethanol Red, of the proteins described above, Prs2 was the only one that presented the same regulation patterns in the two temperature regimes. Prs2 is a member of the ribose-phosphate pyrophosphokinase family and is required for nucleotide, histidine and tryptophan synthesis. The ability to synthetize tryptophan is vital for yeast survival under several environmental stresses. Previous studies have reported that tryptophan uptake at low temperature is a rate-limiting step for *S. cerevisiae* growth. Low temperature increases the rigidity of the plasma membrane, which results in the slower lateral diffusion of membrane proteins, less active membrane-associated enzymes, and a major reduction in membrane transport^26^ Sensitivity of tryptophan uptake at low temperature has been related to a dramatic conformational change in Tat2p^27^. Disorders that affect the stability of the membrane have strong auxotrophic requirements for tryptophan^28,29^. It has been proposed that tryptophan itself provides protection against membrane interruptions. Besides these cell wall/membrane related stresses, it has also been described that tryptophan levels can influence growth recovery after DNA damage^29^. We then speculate that the overexpression of Prs2, in Ethanol Red strain, is mainly related to the requirement of tryptophan and the maintenance of a robust cell membrane to endure thermal stress.

Finally, in the ADY5 experiment, the set of proteins with an identical regulation pattern includes the Hsp82, Mdj1, Ilv6 and Prs2, whose levels were increased, and Gdh1 whose level was diminished. Hsp82 is one of the most highly preserved and synthesized heat shock proteins of eukaryotic cells; while Mdj1 is a mitochondrial Hsp40 involved in many key functions including folding of nascent peptides and cooperation with mt-Hsp70 in mediating the degradation of misfolded proteins^30^. Hsp82 and Mdj1 proteins have a well-known role in heat-response, and their increment has been reported previously by Shui et al. 2015^4^. On the other hand, nothing is known about the role of these two proteins in the cold-response. Furthermore, the presence of two proteins involved in amino-acid metabolism (Ilv6 and Gdh1), in the set of proteins with an identical regulation at sub- and supra-optimal temperatures, was also detected. The first enzyme is an acetolactate synthase, which catalyzes the first step of branched-chain amino acid biosynthesis; and the second is a glutamate dehydrogenase, which participates in the synthesis of glutamate from ammonia and alpha-ketoglutarate^31^. The branched-chain amino acids, such as isoleucine, have already been related to temperature response. Indeed, it has been demonstrated for *Bacillus subtilis* that isoleucine-deficient strains show a cold-sensitive phenotype^31^. In addition, isoleucine has been described as part of a set of amino acids typically abundant in thermophilic organisms, contributing to a high thermal stability of proteins^32^. Likewise, Mara and co-authors observed that overexpression of Gdh1 had detrimental effects on yeast growth at 15 °C^33^. The GDH pathway is involved in the recycling of NADH-NAD by controlling the levels of α-ketoglutarate. The downregulation of Gdh1 may be a strategy to assure NADH-NAD homeostasis, which is essential to ensure an appropriate cellular response to environmental changes.

Taken together, these findings suggest that, in addition to a general thermal response, the strains elicit different protein sets in order to respond to each environmental condition – high or low temperature.

### Comparative analysis of differentially expressed proteins at supra-optimal temperature in the three *Saccharomyces* strains

As in the comparison made of each strain at both temperatures, significant differentially expressed proteins were plotted in a Venn diagram to identify common expression changes in the three strains at 39 °C (Figure 3). The 259 proteins were detected in all strains, however there are not always significant differences in the pattern of expression compared to the respective control temperature (30°C). Only cases where there is a significant increase or decrease will be discussed.

**Figure 3 –.**
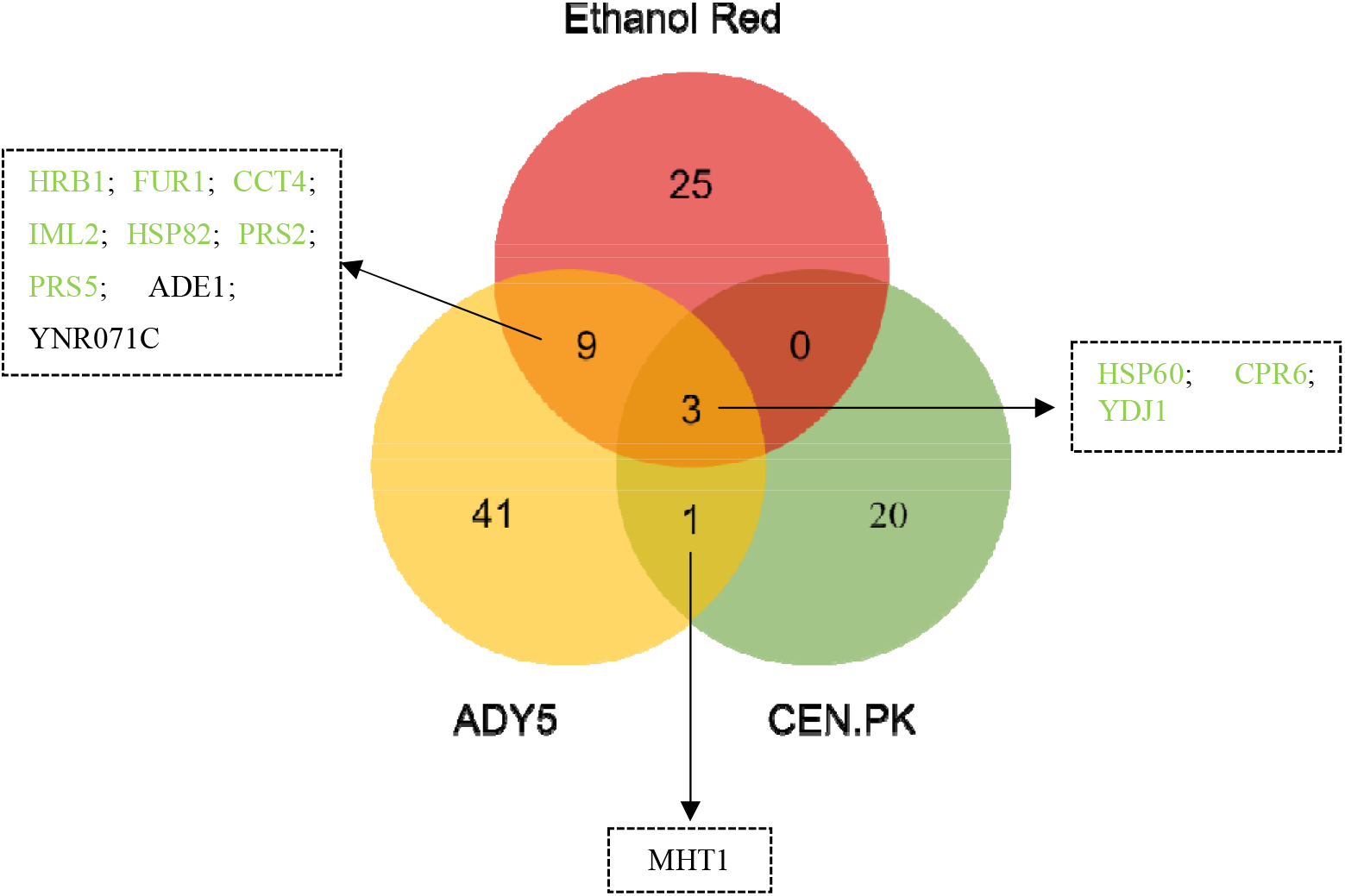
Venn diagram representing the differentially expressed proteins identified in the three *Saccharomyces* strains cultivated at 39°C, in anaerobic chemostat cultures. Proteins are colored according to significant change as follows: green (increased in both strains), or black (decreased in both strains).

Only three proteins were commonly over-represented in the three *Saccharomyces* strains – Hsp60, Cpr6 and Ydj1 – all belonging to the heat shock proteins-family. This result reinforces the view that protein folding is critical for high temperature adaptation and wide-spread among different yeast backgrounds^34^.

In relation to Ethanol Red/CEN.PK, the disparity between the two strains is evident since they do not share any proteins other than those mentioned above. Likewise, ADY5/CEN.PK only have one common protein (Mht1), downregulated in both strains. This homocysteine S-methyltransferase catalyzes the conversion of S-adenosylmethionine (AdoMet) to methionine, regulating the methionine/AdoMet ratio^35^. The repression of this protein allows to keep the amount of methionine reduced, which is essential since methionine suppresses the synthesis of adenylyl sulfate (APS). APS plays an important role in yeast thermotolerance, increasing tolerance to high temperatures^36^

In contrast, the overlap observed between Ethanol Red and ADY5 was larger, with 9 shared proteins (Figure 3). Most of these proteins are related with nucleotide metabolism. Of particular interest is the downregulation of Ade1 and upregulation of Fur1. These two proteins are involved in distinct pathways of nucleotide biosynthesis: the *de novo* pathway and the salvage pathway, respectively. The *de novo* pathway synthesizes nucleotides from amino acids, carbon dioxide and ammonia; whereas the salvage pathway uses preformed nucleosides and nucleobases that are imported or present inside the cell^37^. Our data suggest that at supra optimal temperatures, nucleotide biosynthesis may shift to use the salvage pathway. The *de novo* pathway has a high requirement for energy compared to the salvage pathway^38^. As the cell would be in state of low energy and NADH reserves during temperature adaptation, a reduced demand of these resources could be beneficial. Moreover, the overexpression of Prs2 and Prs5 is also expected since they are key enzymes of 5-phosphoribosyl-1-pyrophosphate (PRPP) synthesis – a compound required for the both *de novo* and salvage synthesis of nucleotides as well as for the synthesis of histidine, tryptophan and NAD^38^. Besides that, both strains show increased expression of proteins related to cellular quality control pathways, namely Hrb1, Cct4 and Iml2^39–41^. The downregulation of a putative epimerase-1 (Ynr071c) was also a common feature between the industrial strains. However, little is known about this protein.

### Detailed analysis of differential proteins at supra-optimal temperature for the Ethanol Red Saccharomyces strain

Ethanol Red is a strain commercially available that has been developed for the industrial ethanol production. Most of the large-scale production of bioethanol is typically carried out at high temperatures (above 35 °C)^42^ and was previously selected as the most thermotolerant^20^. Given that, we decided to analyze more deeply the proteomic alterations of this strain in response to the high temperature.

Our study uncovered some significant elements of the Ethanol Red response to high temperature (Figure 4). The first component involves proteins from the amino acid metabolism. Indeed, we observed repression of proteins involved in arginine biosynthesis, such as Arg1, Arg5, Arg6 and Arg8, and induction of a protein that catalyze the degradation of arginine – ornithine transaminase (Car2). Interestingly, a similar change was recently reported for the thermotolerant yeast *Kluveromyces marxianus* grown at 45 °C in chemostat culture, and therefore could represent an acquired thermotolerance strategy^43^. The reduction of arginine biosynthesis and the acceleration of the conversion of ornithine to proline – catalyzed by the enzyme Car2 ensures both glutamate conservation and proline production. In fact, proline is the main metabolite of arginine metabolism. Under anaerobic conditions, the degradation of arginine leads to intracellular proline accumulation. This increase in proline has been connected with stress protection, being involved in protein and membrane stabilization, lowering the Tm of DNA and scavenging of reactive oxygen species^44^. Moreover, glutamate and proline are precursor metabolites required for the tricarboxylic acid pathway (TCA). It is worth noting that the TCA has been shown not to function completely as a cycle under oxygen deprivation but to be restricted to an oxidative branch leading to 2-oxoglutarate and a reductive branch leading to oxaloacetate^45,46^. Furthermore, the Ethanol Red also increases the expression of several proteins belonging to the nucleotide metabolism.

**Figure 4 –.**
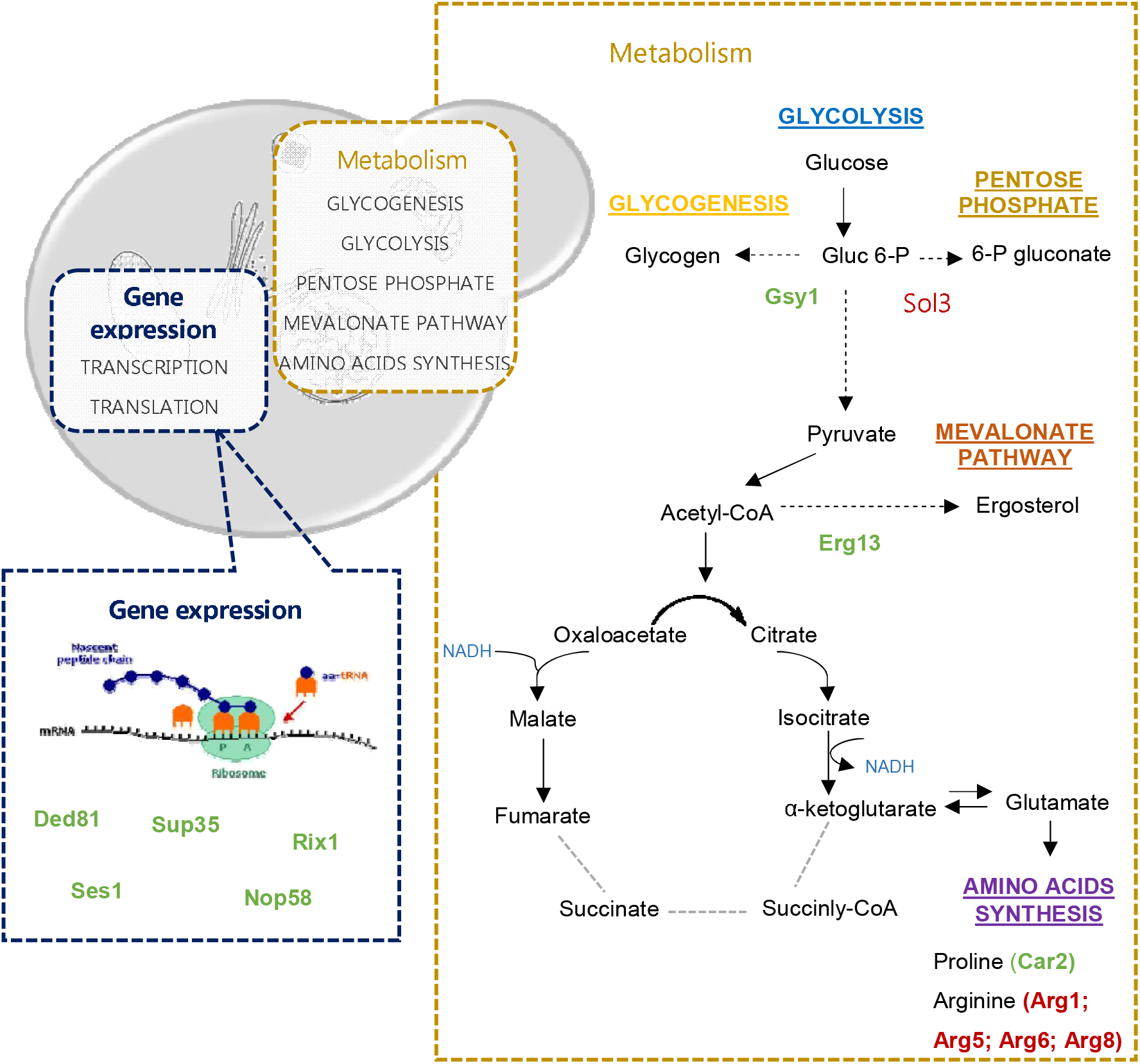
A schematic illustration of the major pathways involved in the unique response of Ethanol Red to supra-optimal growth temperature (39 °C). Under anaerobic conditions, the TCA pathway operates as an oxidative branch and a reductive branch. Broken grey lines represent the point at which the cycle may be interrupted. The precise step of interruption in yeast cells is not yet identified. The proteins are highlighted in green (upregulated) or red (downregulated), indicating their enhancement or repression compared to the levels observed at control temperature (30 °C). In this figure only the proteins are shown which were significantly altered exclusively in the Ethanol Red strain. Altered proteins common to other strains are not shown.

This strain also exhibited a modest downregulation of 6-phosphoglucogalactonase (Sol3) enzyme, which is involved in the so-called oxidative phase in the pentose phosphate pathway (oxPPP). This downregulation at the second step of the pentose phosphate pathway is a key to shifting the glucose metabolism in favor of glycolysis over pentose phosphate pathway. Many of the adaptive responses of yeast to high temperature put a significant additional energy burden on the cells. The increase in temperature causes an increase in the demand for ATP, which can also exert control over the glycolytic flux^47^.

Another notable feature of this strain in relation to the supra-optimal temperature response mechanisms was the overexpression of Erg13 – a protein involved in early ergosterol biosynthesis – and the Gsy1 – a glycogen synthetase. Both proteins were also upregulated at sub-optimal temperature. It has been found that the ability of yeast to tolerate stress is closely related to ergosterol levels. In fact, an increase in the ergosterol content of cell membrane helps to minimize the deleterious effects and maintain a normal membrane permeability^48^. Several studies have reported that the activation of the ergosterol pathways makes yeast cells more resistant/tolerant to a variety of stresses, including low temperature, low-sugar conditions, oxidative stress and ethanol^49,50^. The importance of ergosterol in yeast temperature tolerance has also been demonstrated. It was observed that the compromise of the ergosterol synthesis by mutation in the biosynthesis genes – erg10, erg11, erg19 and erg24 – makes *S. cerevisiae* more sensitive to low temperature^48^. Moreover, Caspeta et al. (2014) also identified Erg3, a desaturase in late biosynthesis of ergosterol, as an efficient target to increase the thermotolerance capacity of yeast^51^. Ergosterol requires molecular oxygen for production and hence it was one of the anaerobic growth factors supplemented to the chemostat medium as it is well known that *S. cerevisiae* has to import sterols under anaerobic conditions^52^. Interestingly, one of the functions attributed to mitochondria under anaerobic conditions is related with sterol uptake^53^. Still, Krantz et al. (2004) also detected upregulation of *ERG13* under anaerobic conditions and concluded that this gene is exquisitely sensitive to addition of even trace amounts of oxygen and that the upregulation could be provoked by addition of water only^54^.

Regarding the increased expression of Gsy1, glycogen has also been reported to be involved with tolerance toward several stresses, such as heat, osmotic and oxidative stress, entry into stationary phase and starvation^55^. The precise role of glycogen in the yeast response to high temperature remains unknown.

In addition, this industrial strain has increased the expression of proteins involved in different translation stages: Ded81 and Ses1 (elongation) and Sup35 (termination), and of proteins related to rRNA processing components (Nop58 and Rix1), which can be a strategy to make the translation process more efficient.

### Comparative analysis of differential proteins at sub-optimal temperature in the three *Saccharomyces* strains

Likewise, the proteomic response of the three *Saccharomyces* strains grown at low temperature was compared (Figure 5). Contrary to that observed at high temperature, no common proteins were found among the three strains, which seems to suggest that a higher disparity between these strains exists in relation to cold-response. However, when directly comparing one strain to another, the number of common proteins is higher than that observed with high temperature.

**Figure 5 –.**
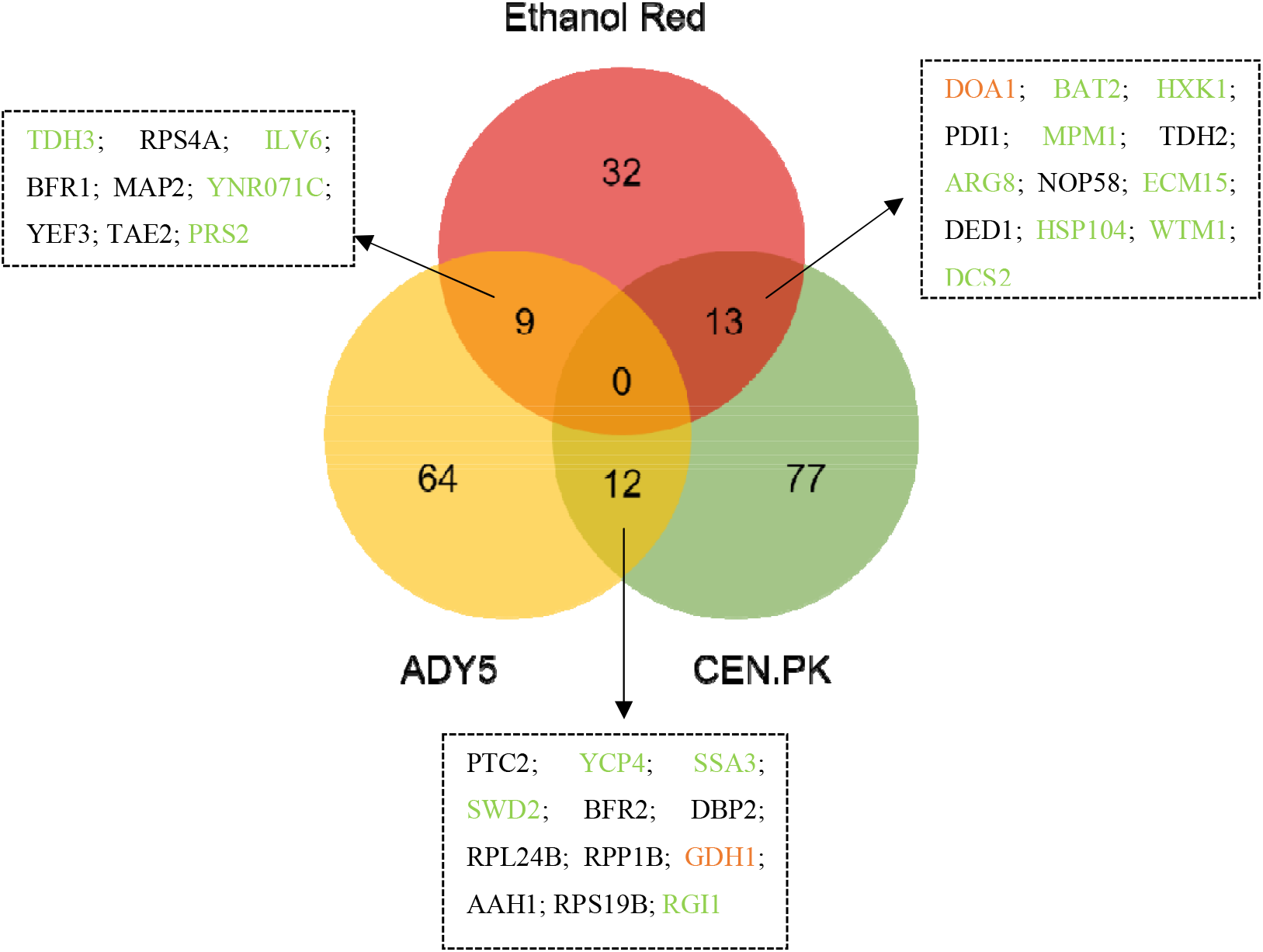
Venn diagram representing the differentially expressed proteins identified in the three *Saccharomyces* strains cultivated at 12 °C, in anaerobic chemostat cultures. Proteins are colored according to significant change as follows: green (increased in both strains), black (decreased in both strains), or orange (opposite expression trends).

### Response of CEN.PK versus Ethanol Red

CEN.PK and Ethanol Red shared 13 proteins, belonging to different functional categories such as protein folding, amino acid and carbohydrate metabolism, transcription and replication, recombination and repair. Although most of these common proteins showed a consistent up-or- downregulation in both strains, differences in the expression levels were observed. Surprisingly, two of these proteins – Ecm15 and Doa1 – exhibited opposite patterns of expression in the two strains. Ecm15 is a non-essential protein of unknown function and is believed to play a role in the biogenesis of the yeast cell wall; while Doa1 is a protein involved in the ubiquitin recycling process, vacuolar degradation pathway and DNA damage repair^56^. Ubiquitin homeostasis is required for the maintenance and growth of the cell. In the case of DNA damage, ubiquitin should be readily available for post-translational modification of proteins involved in the detection, repair and/or tolerance of damage^57^. Therefore, the increased expression of the Doa1 protein in CEN.PK and consequent increase in ubiquitin levels reflects the need for a higher protein turnover, typical of stressed cells. In contrast, the Ethanol Red strain reduces the expression of this ubiquitin protein. Thus, we can interpret that this opposite expression pattern of both strains is related to their different tolerance at low temperatures.

Another key metabolic change common to both laboratory and bioethanol strain lays on carbohydrate metabolism. Both strains increased the expression of hexokinase 1 (Hxk1), which catalyzes the primary step in the glycolytic pathway and reduced the expression of glyceraldehyde-3-phosphate dehydrogenase (Tdh2), which catalyzes the sixth step of glycolysis. Despite being the first step of glycolysis, hexokinase is not the major point of regulation. Glucose 6-phosphate is a branch point between glycolysis, glycogen synthesis and the oxidative arm of the pentose phosphate pathway^58^. The ox-PPP is the main producer of cellular NADPH and is thus critical for the antioxidant defense of cells. It is described that in the presence of oxidants, glucose is channeled towards the ox-PPP through a rearrangement of the carbon distribution. More specifically, the redirection of glycolytic flux through the ox-PPP can be achieved by targeting glycolytic enzymes downstream of Pfk1, such is the case of Tdh2, in which it has already been demonstrated that inhibition leads to a rerouted flux to the PPP, by allowing the accumulation of metabolites upstream of the inhibition point^59^.

Both strains increased the expression of the acetylornithine aminotransferase (Arg8) – a protein involved in the synthesis of arginine. This amino acid is known to have a significant cryoprotective activity, increasing freezing tolerance in yeast cells. The mechanisms underlying this protective property of Arg are not yet known, but it is possible that it acts as an ion coating on the surface of membrane components, avoiding denaturation by the NH2 group in the molecules^60^. Besides, arginine has been shown to reduce cellular oxidative stress, being considered as an effective protectant for proteins, DNA and phospholipids, contributing to maintain intracellular homeostasis^61^.

It is also worth mentioning that CEN.PK had a significant increase in Hsp104 protein at 12 °C although the fold-change was lower in relation to that observed with Ethanol Red. The Hsp104 is considered one of the most crucial thermotolerance-related protein of *S. cerevisiae,* increasing the chances of survival after exposure to extreme heat or high concentrations of ethanol^62,63^. In addition, Hsp104 induction was also observed during late cold response by Schade and his collaborators^9^. It acts as an ATP dependent chaperone that has the distinct ability to solubilize and refold proteins that are already aggregated, allowing other chaperones to have access to otherwise inaccessible surface of the aggregates^64^.

### Response of CEN.PK strain versus ADY5 strain

Likewise, a set of twelve proteins, mainly associated with translation related processes, cell cycle control, nucleotide and amino acid metabolism, were found to be commonly changed in CEN.PK and ADY5 in response to the sub-physiological temperature condition.

To go further, we looked more closely at the common proteins and found three aspects of great interest. First, inspection of the regulation pattern of this set of proteins showed that a glutamate dehydrogenase (Gdh1) exhibited significant changes in both strains, but in opposite directions. Low temperature diminished the concentrations in the wine strain but increased it in the laboratory strain. As mentioned earlier, overexpression of this protein may be detrimental at low temperatures due to a NADH–NAD imbalance^33^. However, a downward shift in the growth temperature also induced an oxidative stress response. It is typically assumed that under anaerobic conditions organisms are spared of the toxic effects of oxidative stress; nevertheless, it has been noted that there are other factors besides oxygen that may induce an oxidative stress response. Gibson et al. (2008) detected an increase in transcription of several antioxidant-encoding genes during small-scale fermentation with *S. cerevisiae* in anaerobic conditions^65^. These authors suggested that the response to oxidative stress may have been triggered by a catabolite repression or the influence of other stresses such as ethanol toxicity. Likewise, we observed an increase in the expression of antioxidant proteins implicated in glutathione synthesis. Gdh1 is involved in glutamate synthesis, which is thought to be part of the defense against oxidative stress through the glutathione system, preventing cold-induced reactive oxygen species (ROS) accumulation^33^. This double and apparently contradictory role of Gdh1 in the cold growth of yeast strains has already been observed by Ballester-Tomás et al. (2015)^66^ and emphasizes the complexity of its responses.

The downregulation of the concentration of Bfr2, which is known for its cold-shock response, was also striking. Bfr2 is involved in protein trafficking to the Golgi and previous studies have demonstrated that its mRNA levels increase rapidly over 5-fold in response to cold shock (when the temperature is changed from 30 °C to 10 °C)^67^. However, this protein may be involved in the recovery from cold shock and the transition back to higher temperatures rather than in survival or adaptation to lower temperatures. This finding underlines that the adaptation to the growth environment and response to shock are distinct mechanisms and straightforward conclusions from one to the other are not always possible.

Another interesting feature is the downregulation of Ptc2 in the CEN.PK strain compared to ADY5, a phosphatase protein known to negatively regulate the high-osmolarity glycerol (HOG) pathway, along with Ptc1 and Ptc3^68^. The HOG pathway is a preserved MAPK cascade in *S. cerevisiae,* which among other functions, is critical for tolerance to different stresses, such as osmotic, citric acid stress and heat stress^69^. Additionally, Panadero et al. (2005) showed that the HOG pathway is clearly activated in response to a downshift in temperature from 30 °C to 12 °C, participating in the transmission of the cold signal and on the regulation of the expression of a subset of cold-induced genes^70^. So, the downregulation of Ptc2, which directly dephosphorylates Hog1 MAPK and limits the maximal activation of the HOG pathway^69^, may be related to this need to activate this pathway.

### Response of ADY5 strain versus Ethanol Red strain

Finally, the comparison of the proteomic profile of the two industrial strains revealed an overlap of 9 proteins, related to the functional categories of carbohydrate metabolism, translation, amino acid and nucleotide metabolism.

One of the most interesting features that has emerged from this comparison is the induction of Prs2. It should be noted that this protein was also identified as significantly altered in the response to supra-optimal temperatures of both strains, Ethanol Red and ADY5 (Figure 3). Therefore, these results suggest that this phosphoribosyl-pyrophosphate synthetase is vital for cellular response to either sub- and supra-physiological temperatures, being essential for the thermotolerance capacity of these two industrial strains.

Apart from this, we found an enzyme of the lower part of glycolysis, Tdh3, which is one of three isoforms of glyceraldehyde-3-phosphate dehydrogenase protein (GAPDH). However, the yeast response does not seem to be so straightforward. For instance, the Ethanol Red strain increased Tdh3 expression, but reduced Tdh2 expression. Likewise, García-Rios et al. (2014)^11^ observed an overexpression of the Tdh3 protein in two wine yeast strains of the genus *Saccharomyces* cultivated at 15 °C. The yeast Tdh2 and Tdh3 GAPDH isoenzymes share extensive sequence homology but appear to have distinct roles in regulating glycolytic flux under oxidative stress conditions^71^.

### Comprehensive proteome analysis of the wine strain at sub-optimal temperature

The ability to ferment at low temperatures (between 10–15 °C) is of paramount importance in the wine industry, since it increases production and retains flavor volatiles, improving the sensory qualities of the wine. In fact, ADY5 strain was previously selected as the best low temperature adapted strain^20^. The main responses observed exclusively in this strain at sub-optimal growth temperature are represented in Figure 6.

**Figure 6 –.**
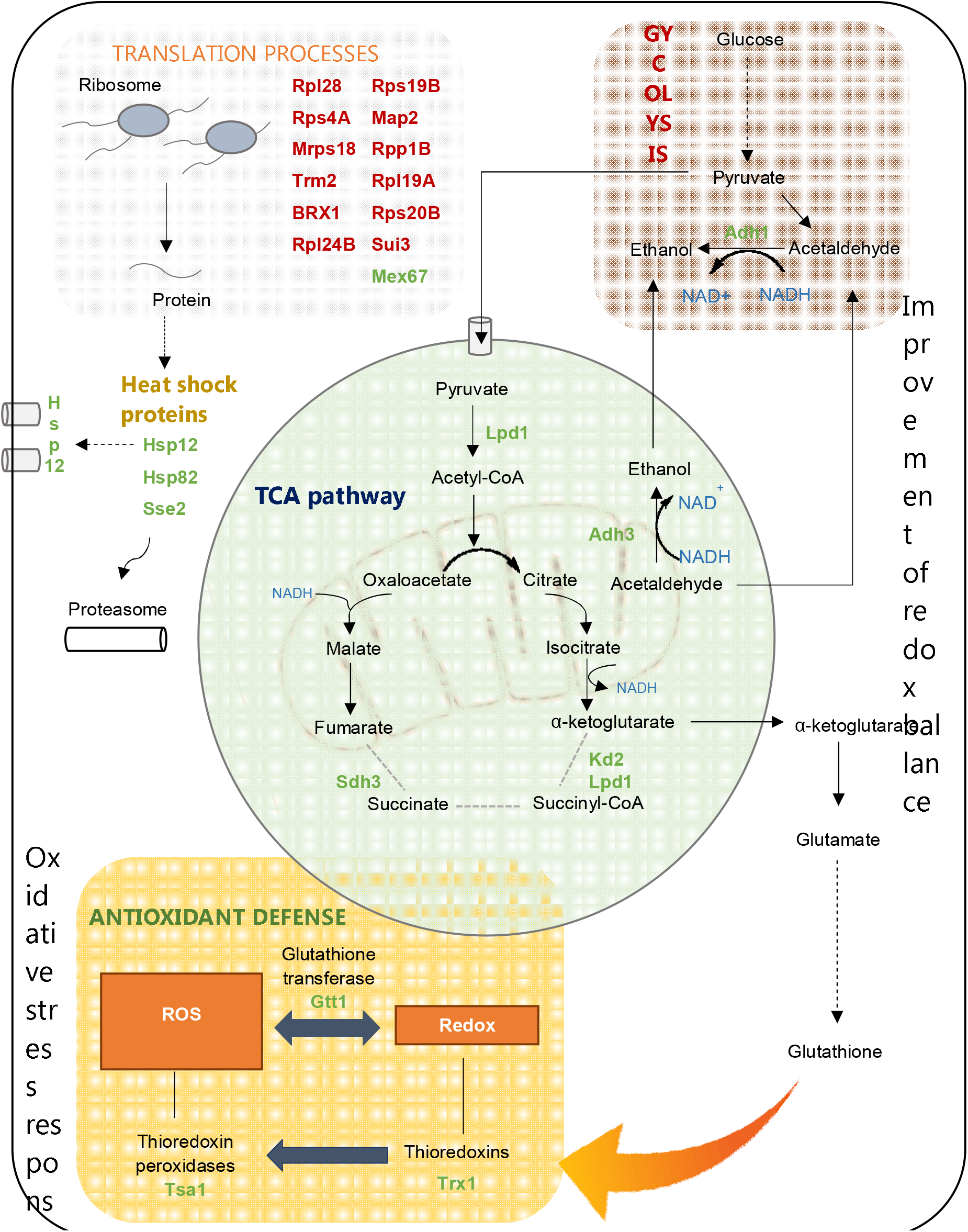
A schematic illustration of the major pathways involved in the unique response of ADY5 to sub-optimal growth temperature (12 °C). Under anaerobic conditions, the TCA pathway operates as an oxidative branch and a reductive branch. Broken grey lines represent the point at which the cycle may be interrupted. The precise step of interruption in yeast cells is not yet identified. The proteins are highlighted in green (upregulated) or red (downregulated), indicating their enhancement or repression compared to the levels observed at control temperature (30 °C). In this figure are represented only the proteins significantly altered exclusively in ADY5 strain. The proteins common to other strains were not represented.

The response of ADY5 to suboptimal temperatures was characterized by a downregulation of several translation-related proteins. For instance, several protein components of the 40S and 60S ribosomal subunit were strongly downregulated^72^. Furthermore, we also noticed the downregulation of translation initiation factors, which influence the translation rate. Yeast adaptation to temperature downshift is dependent of the duration and magnitude of the temperature change^7^. Although several studies have shown an increase in translation-related proteins^7,17,73^ this only happens in the initial and mid phases of cold adaptation. Sahara et al. (2002) observed that after 8 h of exposure to 10 °C, this machinery is significantly downregulated^7^. Similarly, Murata et al. (2005) noticed repression of key genes involved in protein synthesis in yeast cells exposed at 4 °C for 6 to 48 hours^10^. Notwithstanding, this strain presents some interesting molecular mechanisms to overcome the stressors and cell modifications caused by low temperature.

It is worth mentioning the previously reported correlation between low temperature robustness and resistance to oxidative stress in *S. cerevisiae.* Indeed, the fitter strains to fight against oxidative stress were found to display better growth and fermentation performance at low temperature^19^. In the present work, the ADY5 strain increased the expression of a set of proteins belonging to the ROS detoxification systems, including Gtt1 (glutathione transferase), Trx2 (thioredoxin) and Tsa1 (thioredoxin peroxidase). Of these, cytoplasmic thioredoxin 2 is considered one of the most important redox controls together with the glutathione/glutaredoxin system^74^. Considering that these three proteins have only been identified as significantly altered in the ADY5 strain, it can be assumed that they may represent a specific response of the wine strain.

In the category of cell rescue and defense, it is also notable that heat shock proteins are induced, such as Hsp12, Hsp82 and Sse2, which play an equally important role in the response to low temperature in *S. cerevisiae*^75^. Of particular interest was the expression of Hsp12, which has been reported as massively induced in yeast cells under several stress conditions (e.g. heat shock, osmotic stress, oxidative stress or high concentrations of alcohol)^76^. Moreover, Hsp12 is also induced when the cells are exposed to 0 °C^77^ and 4 °C^10^. Hsp12 seems to play a role in cryotolerance of *S. cerevisiae,* presenting similarities with trehalose activity^78^. In fact, Hsp12 acts at the plasma membrane level, protecting it against desiccation and maintaining the integrity of the yeast cells in the freezing stage^75^.

In addition, we also found a remarkable proteomic reprograming in proteins implicated in carbohydrate metabolism and energy production. Of particular interest was the upregulation of two alcohol dehydrogenases, involved in the last step of ethanol synthesis – Adh1 and Adh3. This may represent a strategy of *S. cerevisiae* ADY5 to deal with the redox imbalance produced due to the slower kinetics of alcohol dehydrogenases at low temperature^17^. By increasing the concentration of the main alcohol dehydrogenases, Adh1 and Adh3, the strain overcomes the slower conversion of acetaldehyde into ethanol, avoiding the accumulation of NADH and the blockage of glycolytic flux. Besides, it has been reported that cells of *S. cerevisiae* lacking the Adh3 gene have decreased fitness at low temperature, while its overexpression enhanced cold growth^79^. This may be related with the fact that this protein is part of ethanol-acetaldehyde redox shuttle, being involved in the transfer of redox equivalents from the mitochondria to the cytosol. It is known that low temperatures can compromise the oxidation of mitochondrial NADH, having a strong impact on the intracellular compartmentalization of NAD pools^80^ Therefore, increased expression of proteins like Adh3 may be critical for growth at low temperatures as it helps maintain the optimal balance between the reduced and oxidized forms of nicotinamide adenine dinucleotides.

Furthermore, three proteins from the TCA pathway – Lpd1, Kgd2 and Sdh3 – were also found among the upregulated proteins of this strain. The lipoamide dehydrogenase (Lpd1) is a common component of the multienzyme complexes pyruvate dehydrogenase (PDH), α-ketoglutarate dehydrogenase (OGDH) and glycine decarboxylase (GCV)^81^. Among the several functions of this enzymatic complex, its involvement in one-carbon metabolism is particularly interesting since it is a key pathway that provides single carbon units for the biosynthesis of purines, thymidylates, serine, methionine and N-formylmethionyl tRNA^82^, and part of these components are involved in the cold response. Kgd2 proteins is also a component of OGDH; while Sdh3 is a subunit of succinate dehydrogenase complex, which is supposed to be inactive under anaerobic conditions^46^. Binai et al. (2014) also observed an upregulation of this mitochondrial protein under anaerobic calorie restriction conditions^83^. The upregulation of Sdh3 can be related with the dual function of this protein in the respiratory chain and as a translocase of the TIM22 complex, which is responsible for the insertion of large hydrophobic proteins, typically transporters (carrier proteins), into the inner membrane^84^. The involvement of this complex in yeast response to sub-optimal temperatures has been previously reported by the mutation in Tim18p, a component of the TIM22 complex, which caused a cold-sensitive phenotype^85^.

## Conclusions

Adaptation and tolerance of yeast strains to temperatures beyond an optimum range is crucial for economic and eco-efficient production processes of new and traditional fermentations. Our broad approach highlights the complexity of the proteome remodeling associated with growth temperature perturbation, as well the importance of studying different strains to come to meaningful biological conclusions. The variability of responses of the three strains examined shows that no general rules can be assumed for different *S. cerevisiae* strains, and that the temperature-response is highly dependent of their genetic and environmental background. Overall, proteomic data evidenced that the cold response involves a strong repression of translation-related proteins as well as induction of amino acid metabolism, along with components involved in protein folding and degradation. On the other hand, at high temperature, the amino acid biosynthetic pathways and metabolism represent the main function recruited. Although the changes in the biological processes are, in some cases, similar among strains, differences in the number, type and even in the expression patterns of the proteins were observed and are probably behind the differences in thermotolerance of the strains. Regarding sub-optimal temperature response, the ADY5 strain exhibited a strong activation of antioxidant proteins revealing its importance for low temperature tolerance even under anaerobic conditions. Conversely, at supra-optimal temperatures, Ethanol Red increased the expression of proteins involved in ergosterol and glycogen synthesis, along with Hsp104p, which are known to play a crucial role in heat adaptation.

Our approach and results evidenced the advantages of carrying out a proteomic-wide expression analysis in industrial strains and in an experimental context similar to industrial conditions since it will be valuable for later application in the productive sector. Thus, this study identifies potential proteins involved in yeast thermotolerance that can be useful for the biotechnological industry either through the selection of proper strains or through the development of yeast strains with improved performance at sub- and- supra optimal temperatures.

## Supporting information

Graphical Abstract

## Data availability

The mass spectrometry proteomics data have been deposited to the ProteomeXchange Consortium via the PRIDE^86^ partner repository with the dataset identifier PXD016567.

## Acknowledgments

Financial support is acknowledged to Project ERA-IB “YeastTempTation” (ERA-IB-2-6/0001/2014), and FCT for the strategic funding of UIDB/04469/2020 unit and COMPETE 2020 (POCI-01-0145-FEDER-006684), and BioTecNorte operation (NORTE-01-0145-FEDER-000004). Lallemand Ibéria, SA is acknowledged for the supply of yeast strains. The proteomic analysis was carried out in the SCSIE University of Valencia Proteomics Unit, a member of the ISCIII ProteoRed Proteomics Platform. Authors thank to Luz Valero, researcher of the Proteomic unit, for her valuable support.

## Author Contributions

All authors have been involved in the design of the experiments. KYFL and WVG run the chemostats and took samples for proteomic analysis. TP and LD have analysed the proteomic results and wrote the draft of the paper. All authors read, edited, and approved the final manuscript.

## Additional Information

### Supporting Information

Table S1: Summary of differential proteins found in Ethanol Red, ADY5 and CEN.PK113-7D during sub-optimal or supra-optimal growth temperature experiments.

---

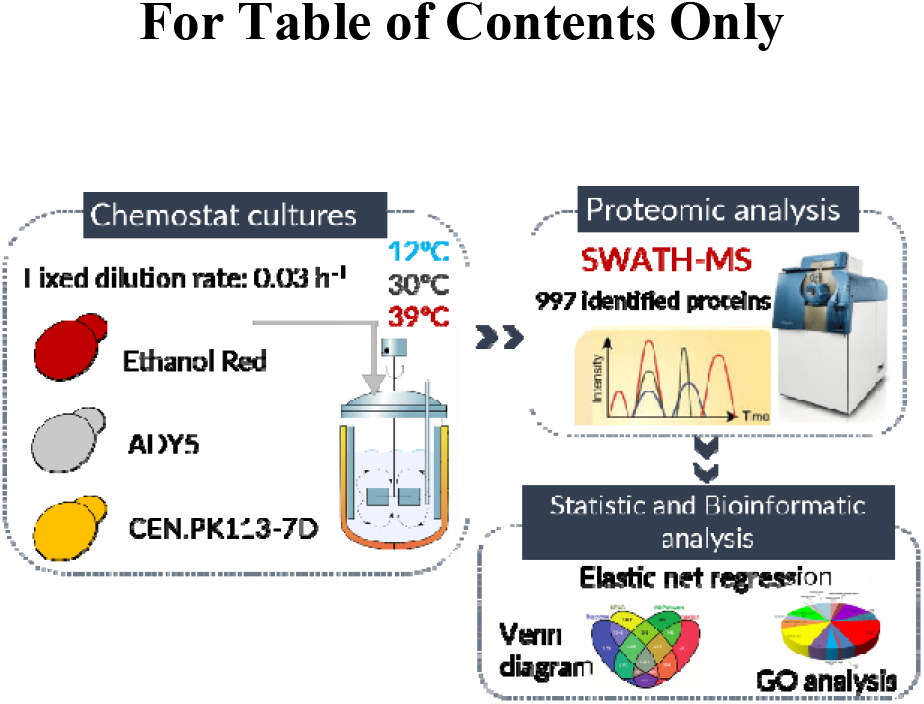

