## Supplementary figures and images for "Differential proteomic analysis by SWATH-MS unravels the most dominant mechanisms underlying yeast adaptation to non-optimal temperatures under anaerobic conditions"

### Graphical Abstract

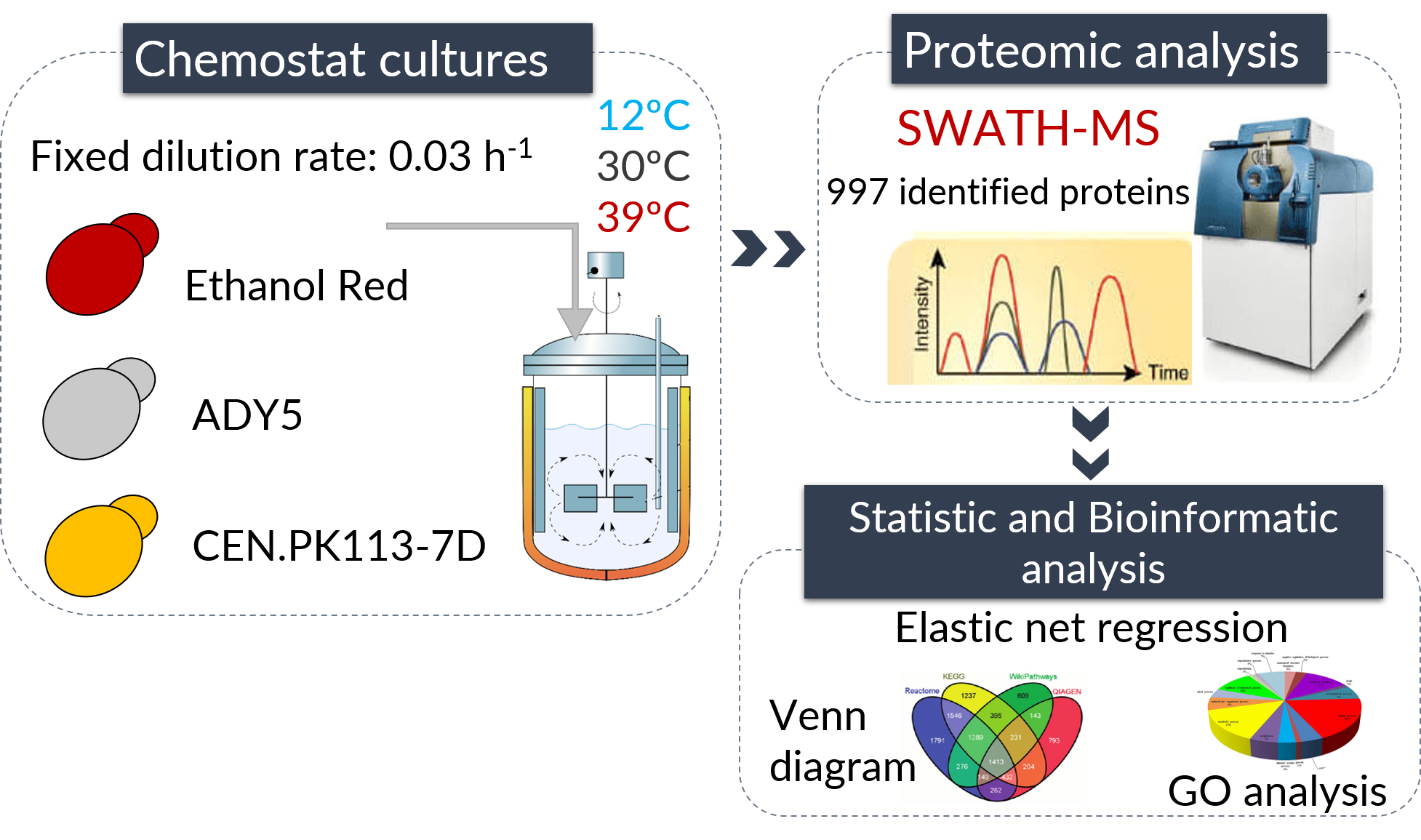
